# Domain dependent orchestrated regulation of bacterial growth, persistence and chemotaxis by an essential GTPase, CgtA, in *Vibrio cholerae*

**DOI:** 10.1101/2021.03.08.434341

**Authors:** Sagarika Das, Partha P. Datta

**Affiliations:** Department of Biological Sciences, Indian Institute of Science Education and Research Kolkata, Mohanpur, Nadia, 741246, West Bengal, India

## Abstract

CgtA, an evolutionarily conserved GTPase, associated with the 50S ribosome controls a broad spectrum of physiological processes in bacteria. It has three structural domains, viz., N-terminal domain (NTD), GTPase domain and C-terminal domain (CTD). CgtA regulates expression of several of genes during nutritional stress in *Vibrio cholerae*. The mechanism of transcriptional regulation by CgtA is unknown, though the NTD concomitantly with the GTPase domain participates in the process. Here, we show that the *in vivo* deletion of the 57 amino acids long CTD of CgtA GTPase of *V. cholerae* is dispensable for viability, contrary to the complete knockdown of *cgtA* gene. Slower growth was observed in *cgtA* knockdown strain with intermittent diauxic lags in minimal media than the CTD deleted strain. Irreversible defect in colony morphology was observed in the cells with CTD deletion. Resuscitation of persister cells occurred when nutritionally deprived complete *cgtA* knockdown cells after growing for longer periods were transferred to nutritionally enriched media. The motility of the *cgtA* knockdown strain was significantly reduced than the wild type cells. Furthermore, CTD deleted cells were also found to be defective in motility, but comparatively lower than *cgtA* knockdown cells. Elongated and slender *V. cholerae* cells were observed by SEM imaging upon *cgtA* depletion, whereas, upon CTD deletion cellular elongation did not occur. Based on our study here, we propose that the CTD of CgtA perceives the nutritional stress response, to which the NTD and GTPase responds.

## 1. Introduction

CgtA GTPase is ubiquitous and conserved between prokaryotes and eukaryotes, where it plays critical role in regulating various vital processes within the cell. CgtA was first identified in *Bacillus subtilis*, where it takes part in the sporulation process of bacteria along with (Trach and Hoch, 1989). other essential functions like ribosome assembly, morphological development and DNA replication. Homologues of CgtA (Obg GTPases) are found in eukaryotes. GTP binding protein 10 (gtpbp10), OBG Protein, in *Xenopus laevis* is involved in early embryogenesis (Jerry et al., 2019). Obg proteins found in plants are involved in chloroplast functioning (Chigri et al., 2009). Another group of Obg family proteins, Drg proteins (developmentally regulated gtp-binding protein) are involved in the mouse brain development (Sazuka et al., 1992). There is a total of ten GTPases in *Vibrio cholerae*, each one of it executing a dedicated set of functions. CgtA GTPase, also known as ObgE GTPase, directs many significant physiological processes within a bacterial cell. The CgtA GTPase is GTP/GDP pool sensor with low affinity for guanine nucleotides and higher exchange rates. (Okamoto and Ochi, 1998; Lin et al., 1999; Wout et al., 2004). Along with various other GTPases found within a cell, CgtA acts in unison in the late stage of 50S ribosome assembly process (Jiang et al., 2006). The participation of CgtA GTPase in the complex process of ribosome assembly makes it a drug-target against pathogenic bacteria. *V. cholerae*, a gram-negative gastrointestinal pathogen, had reportedly caused several pandemic outbreaks in the past. Various antibiotic resistant strains of *V. cholerae* have emerged since then (Das et al., 2020). In cases of severe Cholera, when the oral rehydration therapy may not suffice the prophylaxis, antibiotics need to be administered. Antibiotic resistance of pathogenic strains against the arsenal of drugs available is posing to be a threat against human race.

Designing new drugs against specific drug-targets is critical to combat the issue of antibiotic resistance worldwide. A specific drug target should be an essential protein regulating a cascade of metabolic processes within a cell. Moreover, structurally the drug target should also be unique. Here we present the possibility of CgtA GTPase as a unique candidate as a drug target against antibiotic resistant pathogenic bacteria conferring to its conservation among various bacterial species, essentiality and uniqueness of its structure. The length of CgtA (CgtA_VC_) GTPase in *V. cholerae* O1 biovar El Tor str. N16961 is 390 amino acids long with a molecular weight of 44 KDa. CgtA is structurally unique than other GTPases due to its glycine rich Obg fold with finger-like loops in the N-terminal domain. The highly flexible N-terminal domain acts as a shaft to anchor to the 50S ribosome. Out of the three loops, loop 1 of CgtA_VC_ (spanning amino-acid length ranges from 20-47), and helps to anchor the protein to the 50S ribosome. The loop 2 and loop3 (spanning amino-acid length range from 73-89 and 125-152 respectively) relays the signal of successful interaction of CgtA with the 50S ribosome to the GTPase domain by a flexible leucine rich interdomain linker. Previously, biochemical studies of systematic domain-wise deletion of CgtA had shown its mechanistic aspects (Chatterjee et al., 2019). But the *in vivo* deletion studies are important to know the role of domain-wise functions of CgtA in regulating the intracellular processes. Since the CTD of CgtA varies widely among bacterial species, the outcome of deciphering the role of CTD would be of medical significance. In *V. cholerae*, this can be achieved by *in vivo* deletion of the unstructured57 amino acid length CTD of CgtA_VC_. CgtA is known to be an essential gene in *V. cholerae*. The essentiality of the gene lies in the unsuccessful attempts in deleting the complete *cgtA* gene from the genome of *V. cholerae* unless a complementary copy of wild type gene is supplied. It is not known if the CTD of CgtA_VC_ is essential for its functionality or not. *In vivo* deletion of the CTD of CgtA is necessary to know its essentiality. CgtA GTPase also supports the growth of bacteria. In *V. cholerae*, it was reported that upon CgtA depletion in nutritionally rich LB media, the cells were stuck at the stationary phase for long hours (Raskin et al., 2007). It is not known if the CTD deleted strain would also follow a similar pattern of growth. However, the pattern of *cgtA* depleted *V. cholerae* strain is not yet studied by subjecting the cells to nutritional stress in minimal media. Though, previously it was reported that chemically induced stringent response of *V. cholerae* is similar in many ways to that of *cgtA* depletion condition in LB media. It is not known how the CgtA_VC_ GTPase regulates the general nutritional stress response in *V. cholerae* other than the stringent response. *V. cholerae* well-known to withstand nutrient-poor environments as well as antibiotic stress (Jubair et al., 2012). s. This happens when the cells of *V. cholerae* enter into a state of inactivity/ dormancy for long periods of time. Since, CgtA_VC_ is regarded as a door-keeper of nutritional stress response, then, it might regulate the formation of persister phenotype as well. Understanding the role of CgtA in starvation-induced persister cell formation is necessary since it confers pathogenicity to the *V. cholerae* cells. Furthermore, the dormant persister cells are more tolerant towards antibiotics.

For the cholera pathogen to invade into the gut of the host, chemosensory system plays an important role to sense the fluctuating environmental conditions. *V. cholerae* has three important chemosensory pathways, one of which is known to control its motility (Ortega et al, 2020). Whether CgtA_VC_ has any direct or indirect role in the host pathogenesis can be understood by observing its pattern of motility upon knockdown of entire *cgtA* or deleting its CTD.

Hence, to address some of the above unsolved questions we genetically engineered partial of full *cgtA* deleted strains of *V. cholerae* and designed several biochemical, physiological and analytical experiments. We are depicting results from those experiments below.

## 2. Results

### 2.1. Vibrio cholerae cgtA is essential for cell growth

In order to examine, the effect of CgtA_VC_ in cell-growth, we constructed a *cgtA*-knockdown strain. In this strain, the open reading frame (ORF) of the *cgtA* gene on the chromosome was replaced by homologous recombination with a *kanamycin* resistance gene (kan^r^). This strain maintains pBAD18Cm-cgtA, a helper plasmid containing the *cgtA* gene with its own ribosome binding site and expressed under the control of araBAD promoter. This knockdown strain can be grown in the presence of arabinose. Initially, the primary cultures of both the strains were grown in LB broth for overnight at 37°C. 0.1% arabinose was supplemented in the primary culture of knockdown strain but not for wild-type strain. Both the overnight grown primary cultures were washed to devoid the culture of residual arabinose and were then shifted to the M9 minimal media (supplemented with casamino acids). To completely deplete CgtA from the cells, arabinose was withdrawn from the primary culture of *cgtA* knockdown strain followed by shifting to the M9 secondary culture. It was observed that the cell growth was severely affected as compared to the wild-type strain. The *cgtA* knockdown strain grew for nearly twelve hours before it entered into the stationary phase with intermittent diauxic lags **(FIG.1).** This implies that the CgtA GTPase is necessary to support the cellular growth when subjected to nutritional stress. Thus, the drastically reduced growth rate in the *cgtA* depleted cells appears to be well correlated with the subcellular CgtA concentrations.

**FIG.1.**
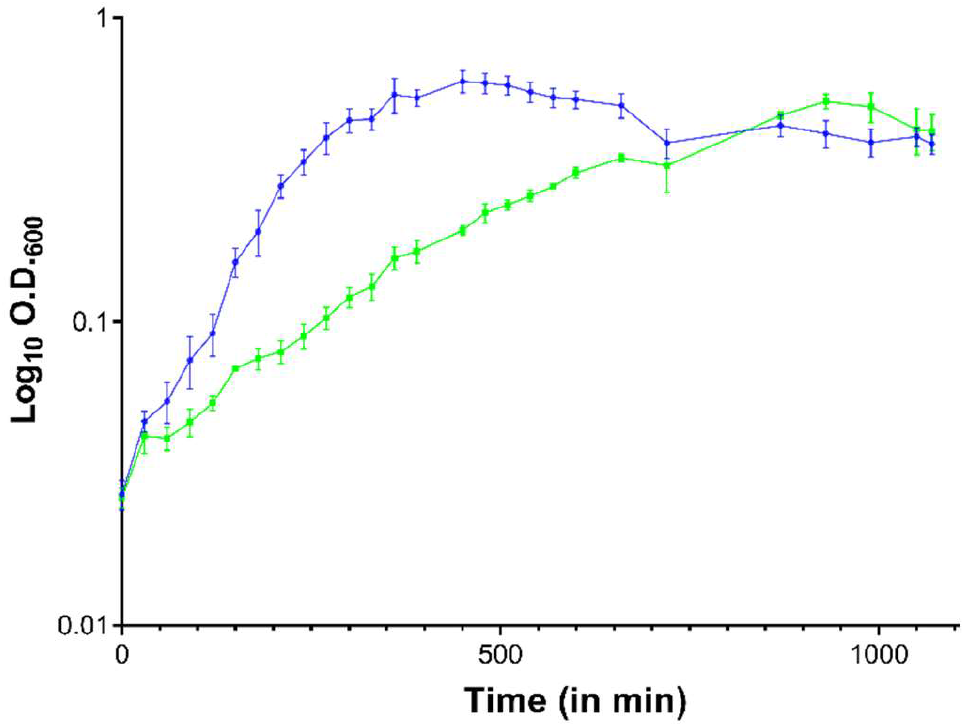
Δ*cgtA::kan^r^* strain when cultured in M9 media supplemented with glucose and casamino acids, exhibits highly retarded growth when compared to the wildtype strain. The knockdown strain starts entering into the stationary phase after 600 minutes of growth.

### 2.2. The C-terminal domain of CgtA_VC_ is not essential for cell viability

The C-terminal domain of CgtA was deleted by replacing the wild-type *cgtA* ORF with Δ*CTD cgtA* allele fused to *kanamycin* resistance gene Δ*CTD cgtA::kan^r^*. Though the helper plasmid pBAD18Cm-cgtA was maintained within the strain, it was no longer needed for the maintenance of viability of the strain. When the freezer stocks of the Δ*CTD cgtA::kan^r^* were streaked on LB agar plates, the strain could grow without the need of arabinose supplementation in the media. On the contrary, the *cgtA* knockdown strain could not be revived from the freezer stocks when streaked on LB agar plate without arabinose supplementation. This implies that the essentiality of the CgtA protein lies in the N-terminal domain and the GTPase domain but not in the C-terminal domain, see **FIG 2.**

**FIG:2.**
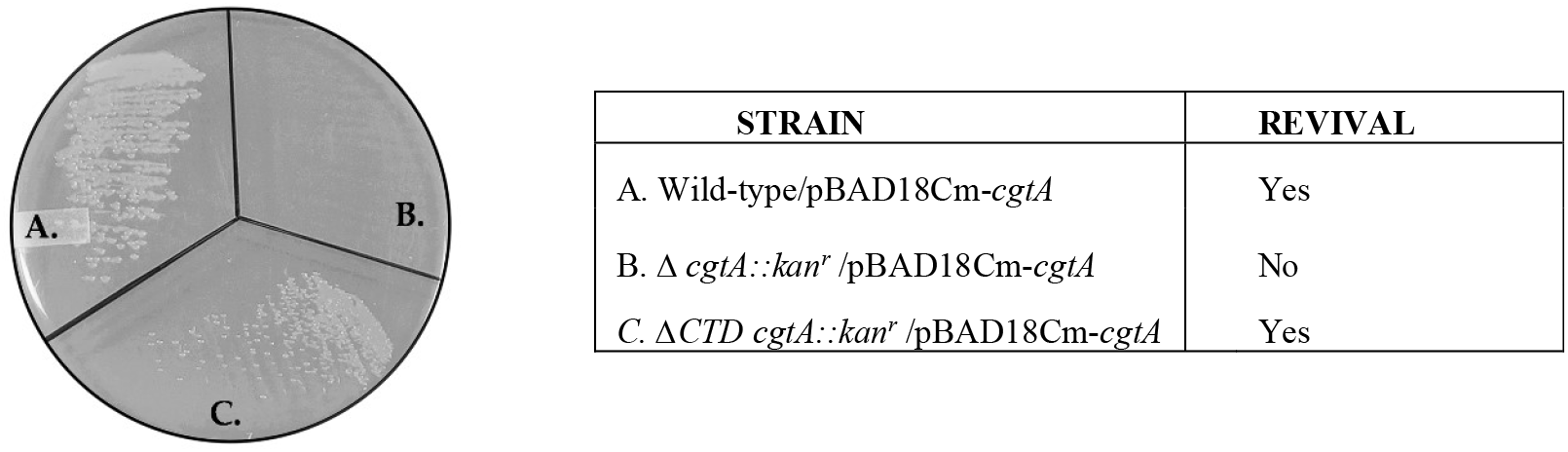
*V. cholerae* strain with *cgtA* gene deletion are unable to revive from glycerol stock when arabinose is not supplemented on LB agar plate. Whereas, *V. cholerae* strain with the C-terminal domain *cgtA* deletion are able to revive without arabinose supplementation indicating the C-terminal domain does not confer essentiality to *V. cholerae*.

### 2.3. Deleting the C-terminal domain of CgtA_VC_ impairs cell growth

Deletion of the C-terminal domain of CgtA impaired cell-growth; but to a lesser extent than the complete *cgtA* knockdown strain. Since, the Δ*CTD cgtA::kan^r^* strain could be revived in the absence of arabinose **(FIG 2)**; hence, arabinose was not supplemented in the growth media for studying the growth pattern. In LB media, the doubling time of the Δ*CTD cgtA::kan^r^* was estimated to be approximately 32 minutes, whereas, for the wild-type strain of *V. cholerae* N16961, the doubling time was estimated to be approximately 19 minutes. The CgtA CTD deletion strain supported bacterial growth better than the complete depletion of CgtA. In M9 media (with glucose and casamino acids supplementation), the growth pattern of wild-type and Δ*CTD cgtA::kan^r^* followed a similar pattern **(FIG 3A)**. Switch point for intermittent diauxic lags were observed in the secondary cultures of both wildtype and Δ*CTD cgtA::kan^r^* in M9 media (with glucose and casamino acids supplementation) after the shift from the primary culture in LB media was observed **(FIG 3B)**. Also, in case of Δ*CTD cgtA::kan^r^* strain, characteristic small colony variants were observed as compared to the wild-type strain implying growth defect. Thus, deleting the CTD elicits nutritional stress in the *V. cholerae* cells in LB media, thus leading to slow growth **(FIG 3C).** This indicates that the C-terminal domain plays an important role in the nutritional stress response upon switching the media from M9 (glucose + casamino acids) to LB.

**FIG: 3.**
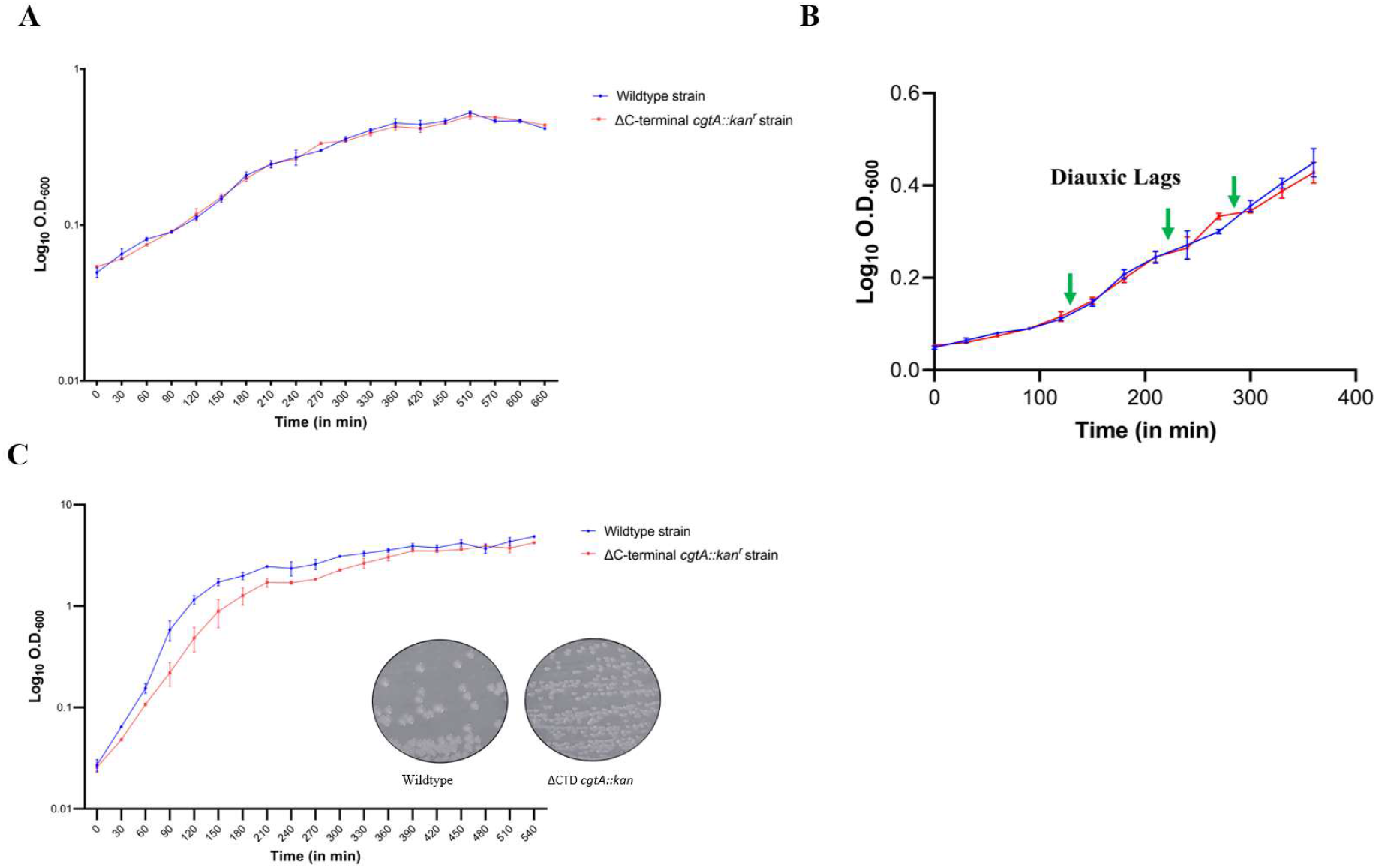
**(A)** *V. cholerae* strain with the CTD deleted CgtA grown in M9 media with glucose and casamino acids supplementation exhibit same growth pattern as wildtype strain. **(B)** Diauxic Lags observed in M9 media (glucose+ casamino acids) for both the strains. **(C)***V. cholerae* strain with the CTD deletion CgtA deletion grown in LB media, grows at a slower rate than the wild type cells. All the experiments are done in replicate and the error bars are the standard error of the mean. This implies, the △CTD *cgtA::kan* is an auxotrophic strain for some nutrients that are deficient in LB media but not in M9+Glucose+casamino acids media. Also, △CTD *cgtA::kan* strain forms pin-hole sized colonies on LB agar plate as compared to wildtype strain indicating slow growth.

### 2.4. △CTD cgtA::kan^r^ is a dominant negative mutation

Even though the deletion of 57 amino-acid long C-terminal domain of CgtA_VC_ is not necessary to support the viability of *V. cholerae*, the colonies grown on LB agar plate at 37°C, overnight-formed pin-hole sized colonies in contrast to the wild-type colonies. When the wild-type cgtA ORF was expressed *in trans*, the growth defect of the Δ*CTD cgtA::kan^r^* cells could not be rescued. The most plausible cause for this observation is that the ΔC-terminal deletion *cgtA* allele is a dominant negative over wild type *cgtA* allele. This implies that, deleting the CTD of CgtA leads to structural changes in the protein that interferes with the function of the wild-type CgtA protein encoded from the helper plasmid. Hence, the constitutive expression of ΔC-terminal deletion CgtA_VC_ mutant protein from the genome, both in the absence and presence (*image not shown*) of *in trans* expression of wild type CgtA_VC_, might result in the constitutive downregulation of genes responsible for bacterial growth defect, resulting in the characteristic small colonies on the LB agar plates, whereas, the colony morphology of *cgtA* knockdown strain was similar to the wild-type strain when grown in presence of 0.1% arabinose. Interestingly, the over-expression of wild-type *cgtA* ORF also leads to characteristic small colonies. This means, the *in vivo* over-expression of CgtA GTPase is toxic for the cellular growth.

### 2.5. CgtA_vc_ contributes to starvation induced persister cell formation in V. cholerae

It is well-known from the earlier studies that, under starvation conditions, *V. cholerae* acquire persister phenotype over a period of time. The persister phenotype of *V. cholerae* cells has epidemiological significance. Sporadic outbreak of Cholera infection occurs when the environmental conditions become favorable for the persister cells to emerge out of the dormant state to proliferate. In our study, the growing culture of all the three strains in M9 minimal media of *V. choleraeN16961in* our study namely wild-type *V. cholerae/* pBAD18Cm-cgtA, Δ*cgtA::kan^r^/pBAD18Cm-cgtA*, Δ*CTD cgtA::kan^r^* were subjected to longer periods of incubation at 37°C for approximately 38 h without arabinose supplementation. With the depletion of the available carbon resources in the growth media, the nutritional stress would aggravate within the cells with time. At the end of the incubation period, the cultures were serially diluted and plated on LB agar media to check for the resuscitation of persister cells. In the CgtA depleted strain, there was a steep reduction in the viable cells as estimated by counting CFU/ml cells after overnight incubation on LB agar media. But the viable cell count was almost similar for wild-type and Δ*CTD cgtA::kan^r^* strains. When, the plates were further incubated till 24 h, micro-colonies of colonies of *V. cholerae* started emerging for the CgtA depleted strain but not for the rest of the strains. The resuscitation of deeply starved *V. cholerae* cells had occurred when regrown on nutritionally enriched media. Thus, depletion of CgtA from the cells elicits a strong starvation induced stress response. Under favorable conditions, the growth of the persister cells resume **(FIG 4).** Thus, the C-terminal domain of the CgtA might not be acting as a regulator of starvation induced persister cell formation though it does trigger nutritional stress as mentioned in the previous section. Either the NTD or the GTPase domain acts as the regulator for persister cell formation. There might be a possibility that the both the NTD and the GTPase domain act simultaneously to regulate the formation of persister cells in *V. cholerae*.

**FIG. 4.**
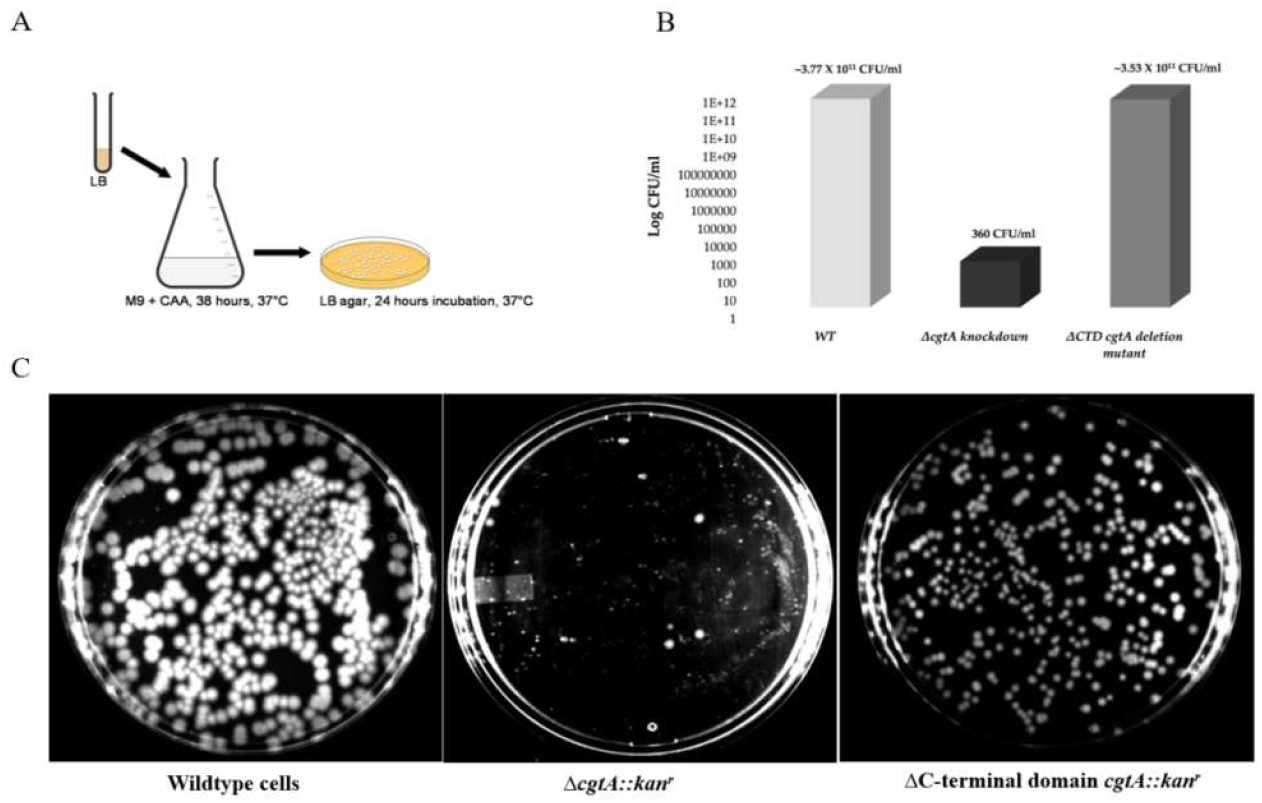
Evidence of persister cell formation **(A)** Workflow for persister cell formation. **(B)** All the strains of *V. cholerae* grown in M9 media with glucose and casamino acids supplementation. CFU/ml was estimated after plating the 38-hour old cultures on LB agar plates and incubated for 12 hours **(C)** The plates were further incubated for next 12 hours and checked for the formation of persister cells. Wild type *V. cholerae* cells and ΔC-terminal domain *cgtA::kan^r^ strain* did not provide any evidence of resuscitation of persister cell formation. Whereas, Δ*cgtA::kan^r^* strain formed tiny cell population on LB agar plate after 24 hours incubation., indicating resuscitation of persister cells.

### 2.6. CgtA regulates the chemotactic behavior in V. cholerae

The RNA-Seq experiment revealed that a set of nineteen genes involved in chemotaxis was downregulated in CgtA depleted *V. cholerae* strain. Out of the nineteen differentially regulated genes, most of them fall into the category of Cluster II/F6 chemosensory system of *V. cholerae*. Out of the three major chemosensory systems of *V. cholerae*, Cluster II/F6 is only responsible for regulating the chemotactic behavior (Ortega et al, 2020). Hence, to regulate the chemotaxis, *cgtA* either plays a direct or indirect role in the differential regulation of the chemotactic genes. To find out which domains of CgtA are highly involved in the role of chemotaxis, we performed motility swarm assays in LB soft agar. In the order of decreased order of motility, the wild-type strain was highly motile, followed by Δ*CTD cgtA::kan^r^* whereas; the CgtA depleted strain was the least motile **(FIG 5A).** Furthermore, many chemotaxis related genes were found to be downregulated through RNA-Seq experiment in Δ*cgtA::kan^r^* strain as depicted through heat map **(FIG 5B)**, but not in the case of Δ*CTD cgtA::kan^r^* strain. Thus, the motility of *V. cholerae* cells is severely affected only if the NTD and the GTPase domains are deleted along with the CTD of CgtA. Since, the motility is less affected in Δ*CTD cgtA::kan^r^* strain; this implies that the chemotactic gene regulatory roles are mostly under the control of NTD and GTPase domain either independently or combinatorially. Moreover, the swarming ability of all the three strains of *V. cholerae* is in concurrence with their respective growth rates. The chemotaxis related genes downregulated in the Δ*cgtA::kan^r^* strain is enlisted in **Table 1**.

**FIG.5.**
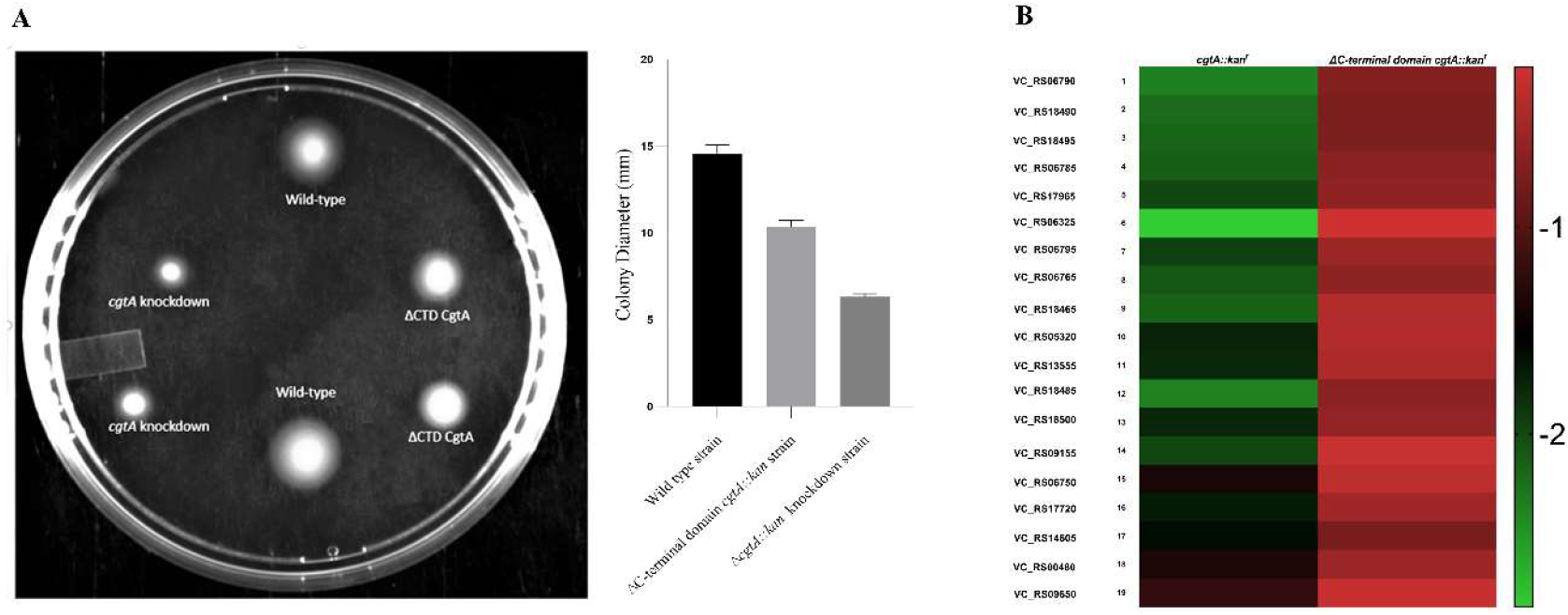
Analysis of motility by swarm-plate assay. (A) The *cgtA* knockdown *V. cholerae* strain exhibits defect in motility to a larger extent followed by the ΔC-terminal domain deletion of CgtA *V. cholerae* strain. (B) All the 19 chemotaxis related genes differentially expressed are down-regulated in Δ*cgtA::kan^r^ V. cholerae* strain. The Log_2_ Fold change is depicted by the heat maps. The genes involved in chemotaxis are not downregulated in △C-terminal domain *cgtA::kan^r^ strain*, indicating the less motility might be linked due to slow growth.

**Table 1:**
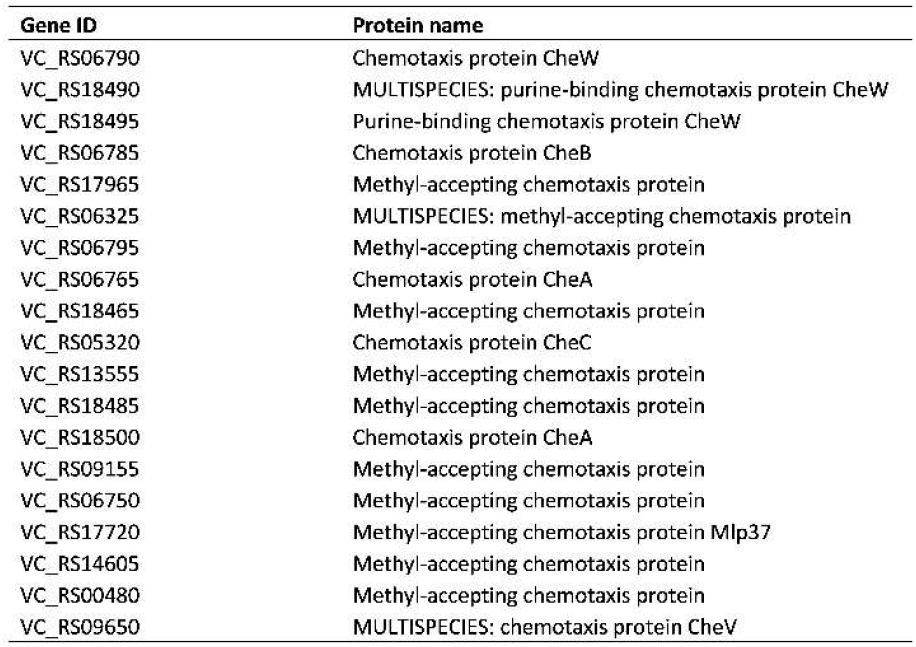
List of 19 chemotaxis related genes downregulated in *cgtA::kan^r^* strain

### 2.7. The C-terminal domain of CgtA might act a sensor for nutritional stress

In order to understand the role of the CTD, the growth as well as the morphology of the wild-type and CTD deleted *cgtA* strains of *V. cholerae* was examined by growing the strains in LB as well as M9 minimal media supplemented with glucose and casamino acids. As aforementioned, in the LB media, the strain with CTD deleted *cgtA* was found to be less than the wild-type strain **(FIG 3C)** Increment in doubling time for the CTD *cgtA* deleted strain was observed in all the experimental replicate considered for the study, thus hinting at the role of CTD in growth in LB media. Whereas, in M9 media supplemented with glucose and casamino acids, the growth pattern of the CTD deleted *cgtA* mutant strain was similar to the wild-type strain (**FIG. 3B)**. The slow growth observed in the exponential phase in LB media disappeared in M9 media with glucose and casamino acids. Thus, the LB media was nutritionally deficient (either for carbon sources or free metabolizable amino acids), for the CTD deleted *cgtA* strain.

Supporting the above fact, when scanning electron microscopy (SEM) imaging (**FIG. 6)**. was performed for all the 3 strains (namely wild-type, *cgtA* knockdown and CTD *cgtA* deleted strains) in both types of media. Morphologically, all the three strains appeared similar when the exponentially growing cells were considered for imaging. Surprisingly, when all the 3 strains were grown in LB media, the *cgtA* knockdown strain from the exponential phase appeared exceptionally elongated and stressed as compared to the wild-type cells. Also, an abundance of small sized cell population was observed in CTD deleted *cgtA* strain, indicating delayed growth. Thus, the CTD might be acting as a nutritional stress sensor linked to growth, whereas, the N-terminal Obg domain and the GTPase domains act either in unison or separately to respond to the stress signal (stress-responders). Also, this study establishes that CgtA depleted strain is an auxotrophic mutant either for carbon or any amino acid. As, LB media contains mixture of oligopeptides in contrast to free amino acids in casamino acids. CgtA knockdown strain being a metabolically weakened strain might not be able to utilize the oligopeptides present in LB and hence its growth is supported when casamino acids are added to the M9 media. Also, it is known that grown in LB media is carbon limited (Sezonov G. et al., 2007). Thus, glucose deprivation linked stress could also be the cause of the above observations.

**FIG.6.**
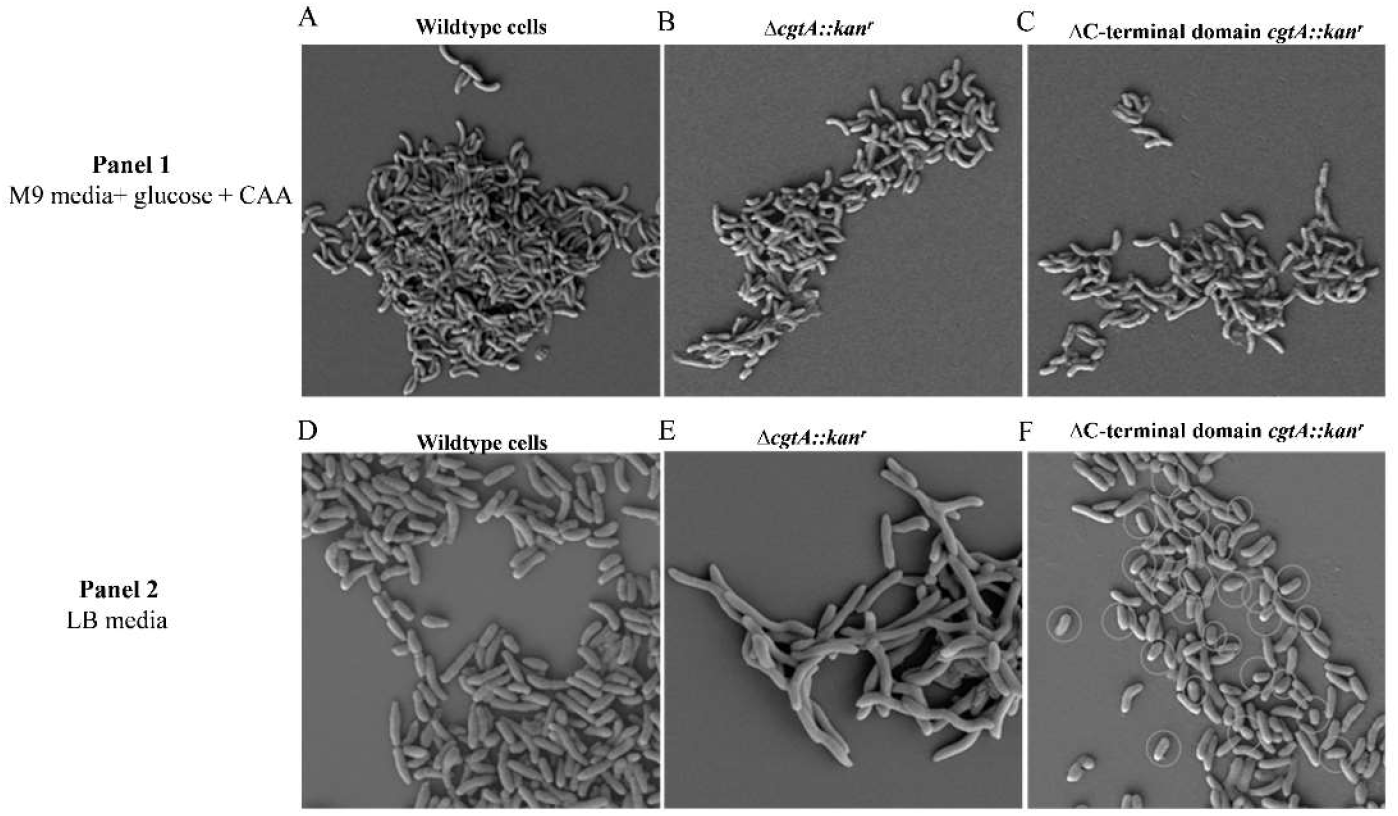
Scanning Electron Microscopy Imaging **Panel 1**: Exponentially growing cells (A, B, C) grown in M9 media with glucose and casamino acids supplementation show no distinct morphological difference (Magnification 5.00 KX). **Panel 2**: Exponentially growing cells grown in LB media (D) Wild type *V. cholerae* cells (E) The *cgtA* knockdown *V. cholerae* strain becomes highly elongated (F) △C-terminal domain *cgtA::kan^r^ strain* with abundance of small size cell population, indicating slow growth. (Magnification ~10.00 KX).

### 2.8. Results from surveying the global transcriptional profile of the CgtA depletion and △CTD cgtA::kan^r^ strains in M9 minimal media with glucose and casamino acids supplementation

In order to understand the underlying physiology of *V. cholerae* when the intracellular CgtA is completely depleted or the CTD of CgtA is deleted, the transcriptome of all the strains were investigated. The strains were grown in M9 minimal media (with glucose and casamino acids supplementation). Since, CgtA tracks the nutritional status of the cell; growing in LB was proving to exert nutritional stress to the *cgtA* knockdown strain. We were interested to see the general stress response effects exhibited by CgtA other than nutritional stress response. To understand the transcriptional landscape of the CgtA regulated genes, we carried out RNA-Seq studies. Also, the general stress response proteins regulated by CgtA GTPase also became evident. The total RNA from all the strains were isolated at low OD_600_ (0.2-0.3) followed by cDNA library preparation, adaptor ligation and Illumina platform based Next Generation Sequencing (NGS). **(FIG 7.)** The number of mapped cDNA reads corresponded to a total of 3,657 genes on both chromosome I and chromosome II of *V. cholerae* N16961. For all the strains, greater than 98% of the reads were successfully mapped. A total of three comparative transcriptional profiling studies were performed. In the first category, when the CgtA depletion strain was compared with the wild-type strain, 256 genes were differentially expressed. Out of which, 226 genes were downregulated and 26 were upregulated. **(FIG 8A).** On the contrary, Δ*CTD cgtA::kan^r^* showed negligible differentially regulated gene pattern **(FIG 8B).** The gene ontology (GO) analysis revealed that a wide variety of molecular functions and biological processes were affected due to CgtA depletion. **(FIG 9)**

**FIG.7.**
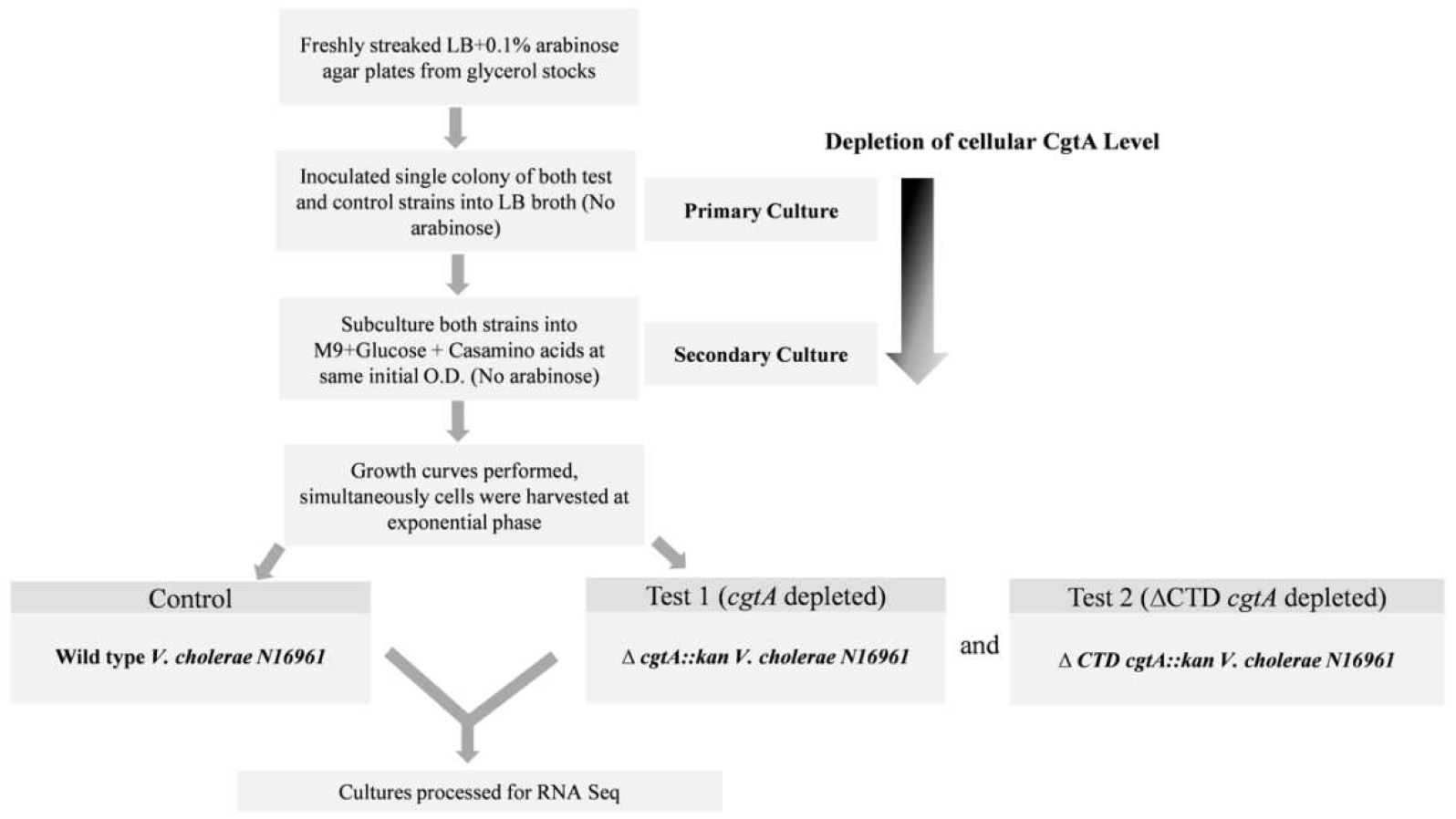
Workflow for the preparation of samples for transcriptional profiling by RNA-Seq

**FIG. 8.**
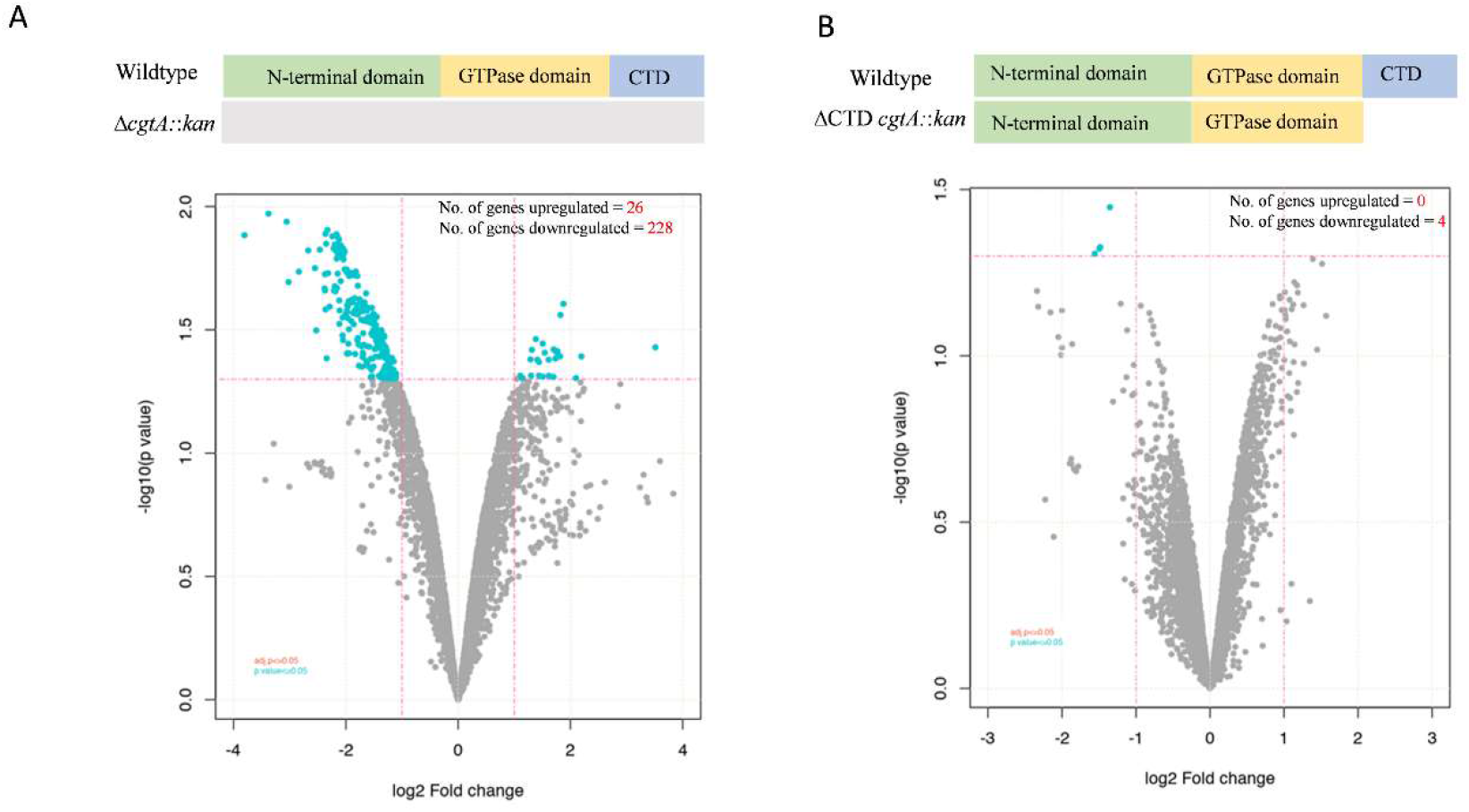
Analysis of differentially regulated genes in △*cgtA::kan^r^* and △C-terminal domain *cgtA::kan^r^* strains in comparison with wildtype *V. cholerae*. **(A)** The volcano plot depicts that a total of 254 genes are differentially expressed in *cgtA* knockdown strain. **(B)** The volcano plot depicts that upon C-terminal domain deletion of CgtA only 4 genes are downregulated. Thus, C-terminal has negligible role in gene regulatory functions. The regulatory functions are majorly carried out by the N-terminal Obg domain and GTPase domains.

**FIG.9.**
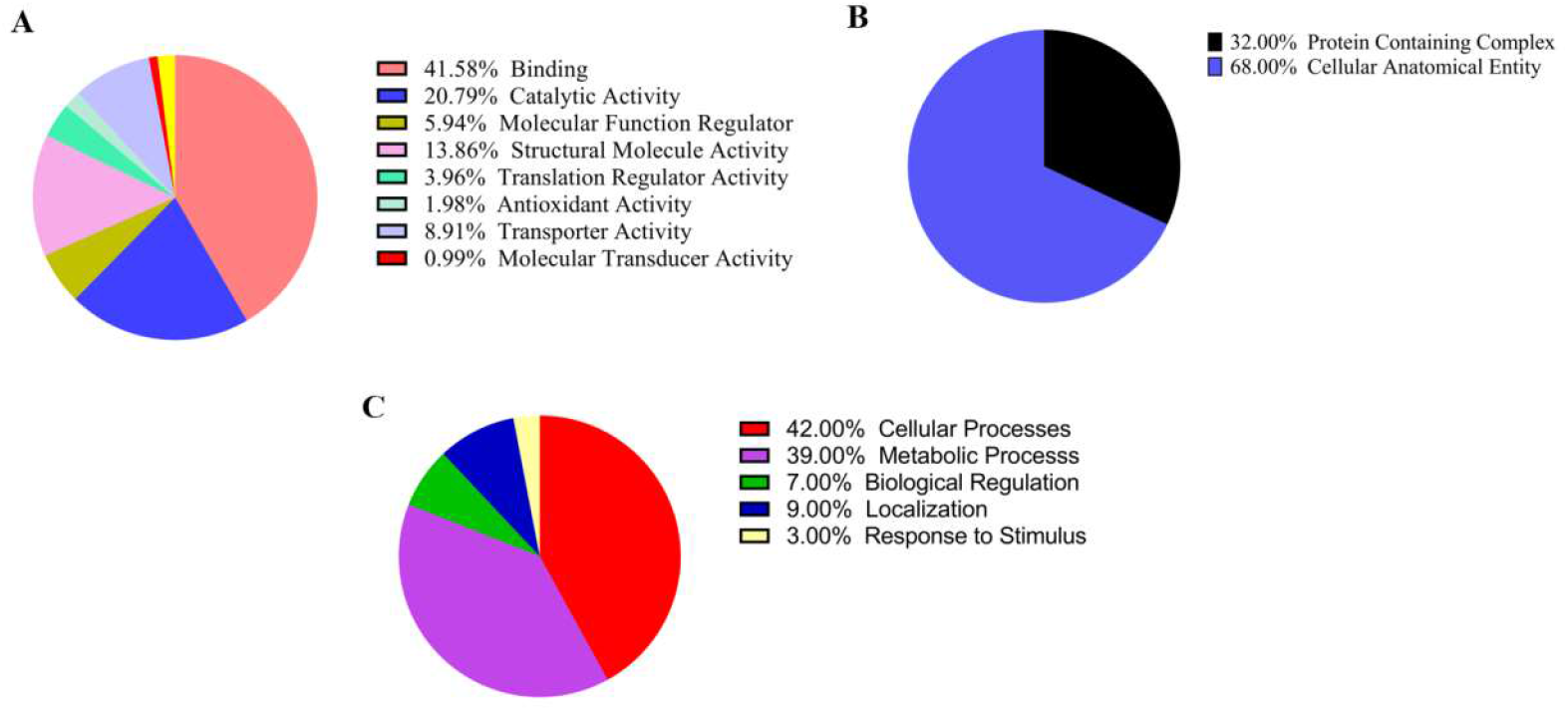
Gene Ontology Analysis for functional characterization of CgtA in *V. cholerae* **(A)** Molecular Function **(B)** Cellular Components **(C)** Biological Process

It is evident from the RNA-Seq data that, the differential expression pattern of the genes is mainly carried out by the N-terminal domain and the GTPase domain individually or due to the interaction of the two domains, but definitely not due to the C-terminal domain. A large number of genes of metabolic interest were affected upon CgtA depletion. Thus, it implies that the transcriptional regulatory functions are dependent upon N-terminal Obg domain and GTPase domain.

A complex gene regulatory network of *cgtA* is depicted in **FIG 10.**

**FIG.10.**
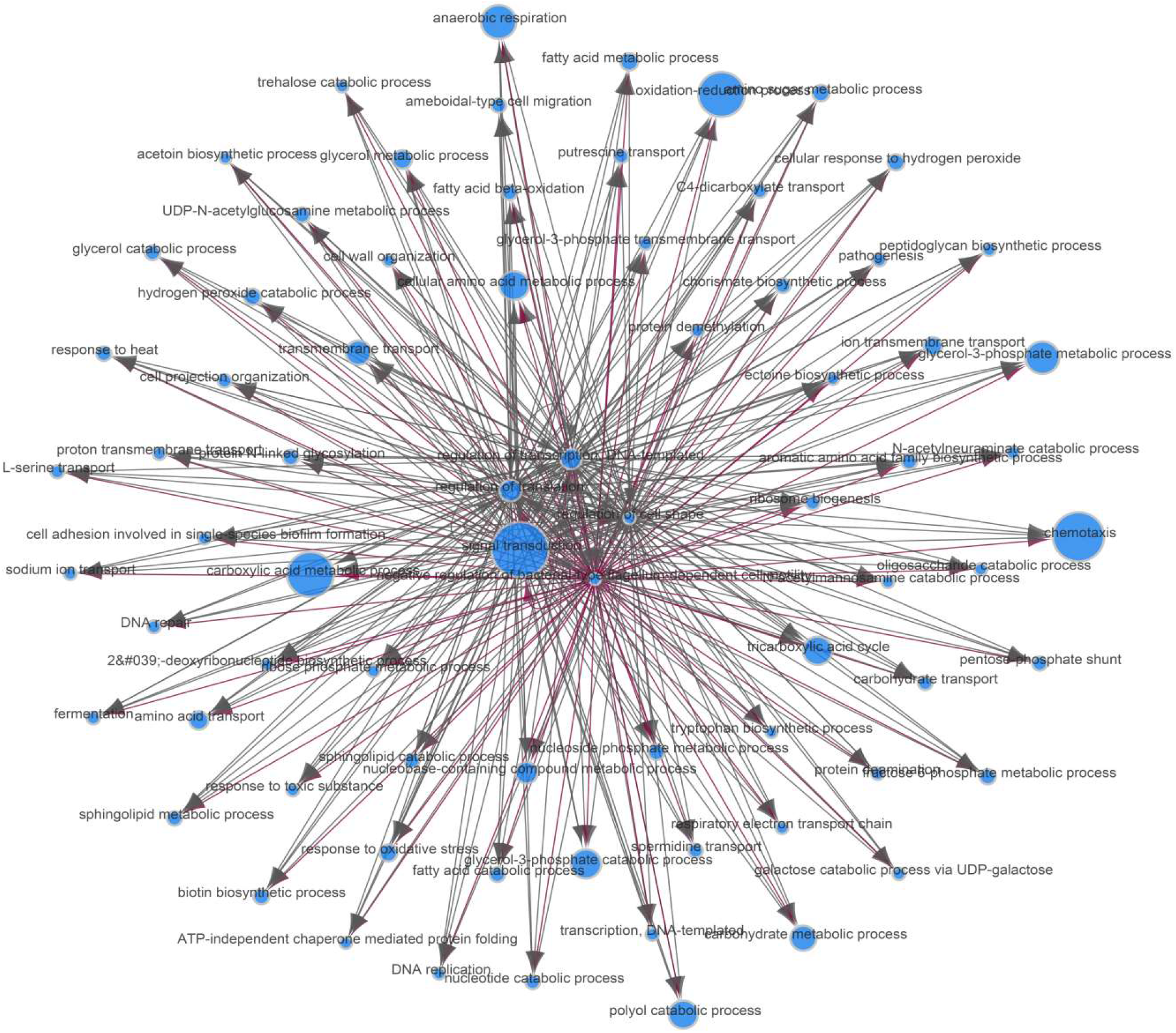
Slow growth is a consequence of low metabolism as evident from “Gene Ontology Network” of downregulated genes in △*cgtA::kan V. cholerae* strain

### 2.9. Analyzing the transfer RNA (tRNA) abundance in cgtA knockdown condition

Since CgtA is a stress response protein, it is expected that a variety of tRNA genes would be differentially expressed. During stressful conditions, cells alter their tRNA abundance to selectively regulate translation of proteins (Torrent M et al., 2018). The pattern of tRNA expression during *cgtA* knockdown condition was evident from RNA-Seq data **(FIG 11).** Many tRNA genes were down-regulated in this condition. This might be due to the rearrangement of tRNA pool due to the general stress response caused due to *cgtA* depletion in *V. cholerae* cells in order to influence protein synthesis.

**FIG.11.**
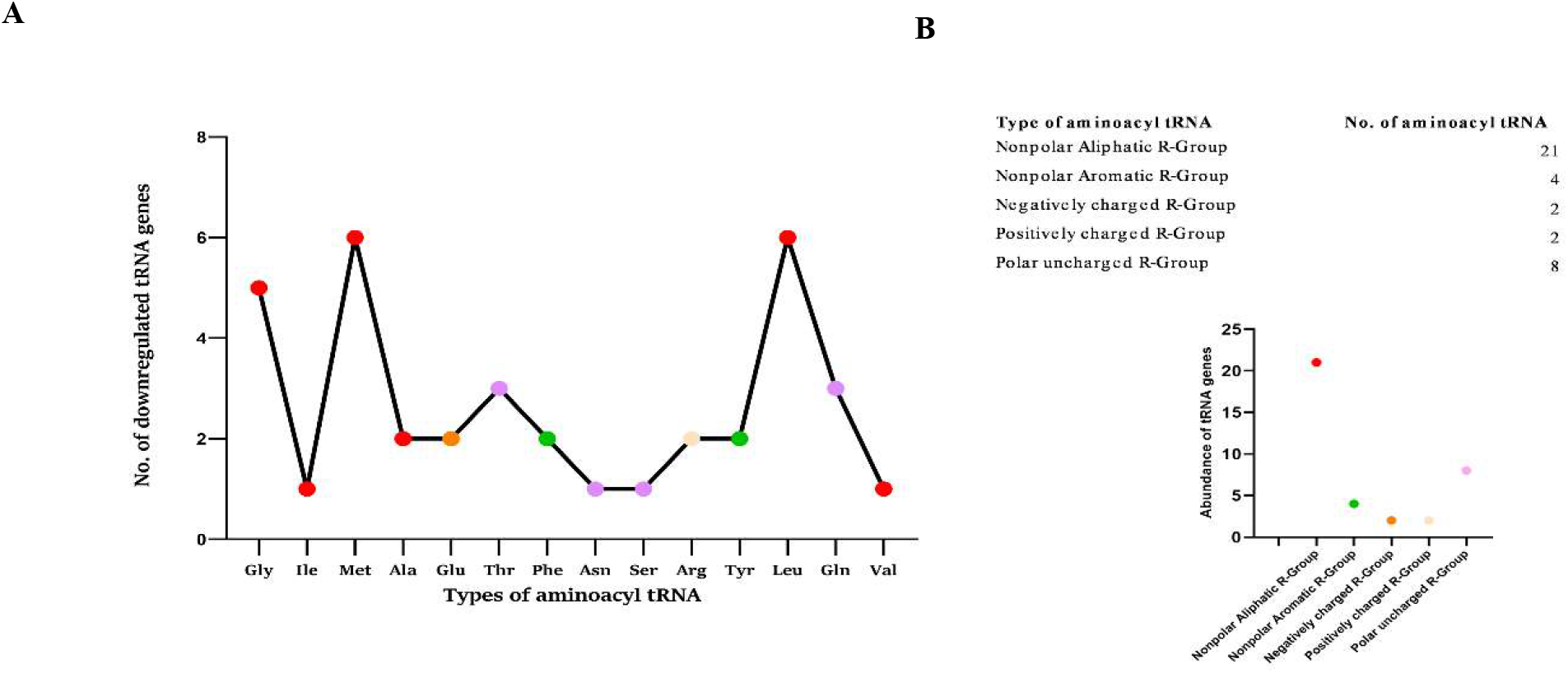
**(A)** Analysis of tRNA genes downregulated in △*cgtA::kan^r^ V. cholerae*. **(B)** Differentially regulated aminoacyl tRNA genes grouped according to the properties of the side chain of the amino acids.

## 4. Experimental Procedures

### 3.1 Molecular biology methods

PCR primers were designed manually and analyzed using the web-based OligoAnalyzer tool by Integrated DNA Technologies (IDT) and manufactured by IDT and GCC Biotech at 25 nmole scale. PCR amplification of various DNA fragments was performed using Eppendorf Mastercycler nexus X2. For overlap extension PCR, NEB Q5 high-fidelity DNA polymerase was used. For A-tailing for TA cloning, NEB Taq Polymerase was used. Agarose gel electrophoresis, restriction enzyme digestions and ligations were performed using standard techniques. Extractions and purifications were performed according to the manufacturer’s instructions in kits purchased from QIAGEN.

### 3.2 Bacterial strains

Streptomycin resistant *V. cholerae N16961* was the target for the construction of the mutant strains. pKAS32 suicide vector-based constructs, which requires Pi protein for replication, was maintained in *E. coli* SM10λpir. Plasmids based on pBAD18-Cm expression vector were maintained in *E. coli* DH5a. *V. cholerae El tor N16961* was used as a recipient strain. To facilitate bipartite mating *E. coliSM10λ pir* was used as a donor strain. *E. coli DH5α* as well as *E. coli SM 10λ*pir were used as a cloning strain. Cells were cultured in lysogeny broth (LB) medium or solidified medium (1.5% agar) at 37°C. High concentration of streptomycin antibiotic was incorporated into LB agar media to facilitate counter-selection process. TCBS (Thiosulfate-citrate-bile salts-sucrose agar) selective agar media (Himedia) was also used to select the *Vibrio cholerae* cells after biparental mating. Bacterial cultures were stored in LB broth with 25% glycerol at −80°C, and grown in LB broth and LB agar. Ampicillin at a concentration of 100μg/ml and 150μg/ml for *E. coli* and *V. cholerae* respectively; Streptomycin at a concentration 100 μg/ml-2mg/ml for *V. cholerae*, Kanamycin at a concentration of 40μg/ml for *V. cholerae* and Chloramphenicol at a of 34μg/ml and 3μg/ml for *E. coli* and *V. cholerae* respectively were added to the media.

### 3.3 Plasmids for creating the in vivo mutant strains of V. cholerae

The suicide plasmid used to create the *in vivo* mutant strains, pKAS32, contains the ampicillin marker gene, which allows for the selection of antibiotic resistance. pKAS32 plasmid also contains the *rpsL* gene of *E. coli*, the locus that encodes ribosomal protein S12, which is useful as a positive selection marker from the fact that mutations in *rpsL* gene that confer streptomycin resistance (Sm^r^) are recessive in a bacterial strain that expresses wild-type *rpsL* protein. The *E. coli rpsL* expressed frompKAS32 is dominant over the *rpsL* expressed from *V. cholerae* genome that aids in counter selection when plated on high concentration of streptomycin. This will lead to the selection of colonies that have undergone an allelic exchange by homologous recombination. Also, pKAS32 contains the origin of replication R6K which ensures that the plasmid replication is dependent upon π, the *pir* gene product. In this study, an extra XbaI restriction site was introduced at the MCS of pKAS32. pKAS32-*UK^r^D* was created by digesting by pGEMT-*UK^r^D* with XbaI and ligating the 2-Kb insert *UK^r^D* with similarly digested pKAS32 with T4 DNA ligase. By using a similar strategy, the construct for introducing the C-terminal deletion *cgtA* allele was also created. Competent *E. coli SM10 λ pir* were selected with ampicillin. Success of the ligation reaction was verified by colony PCR, plasmid extraction, digestion, visualization of a 2Kb band after gel electrophoresis and DNA sequencing.

**Table 2:**
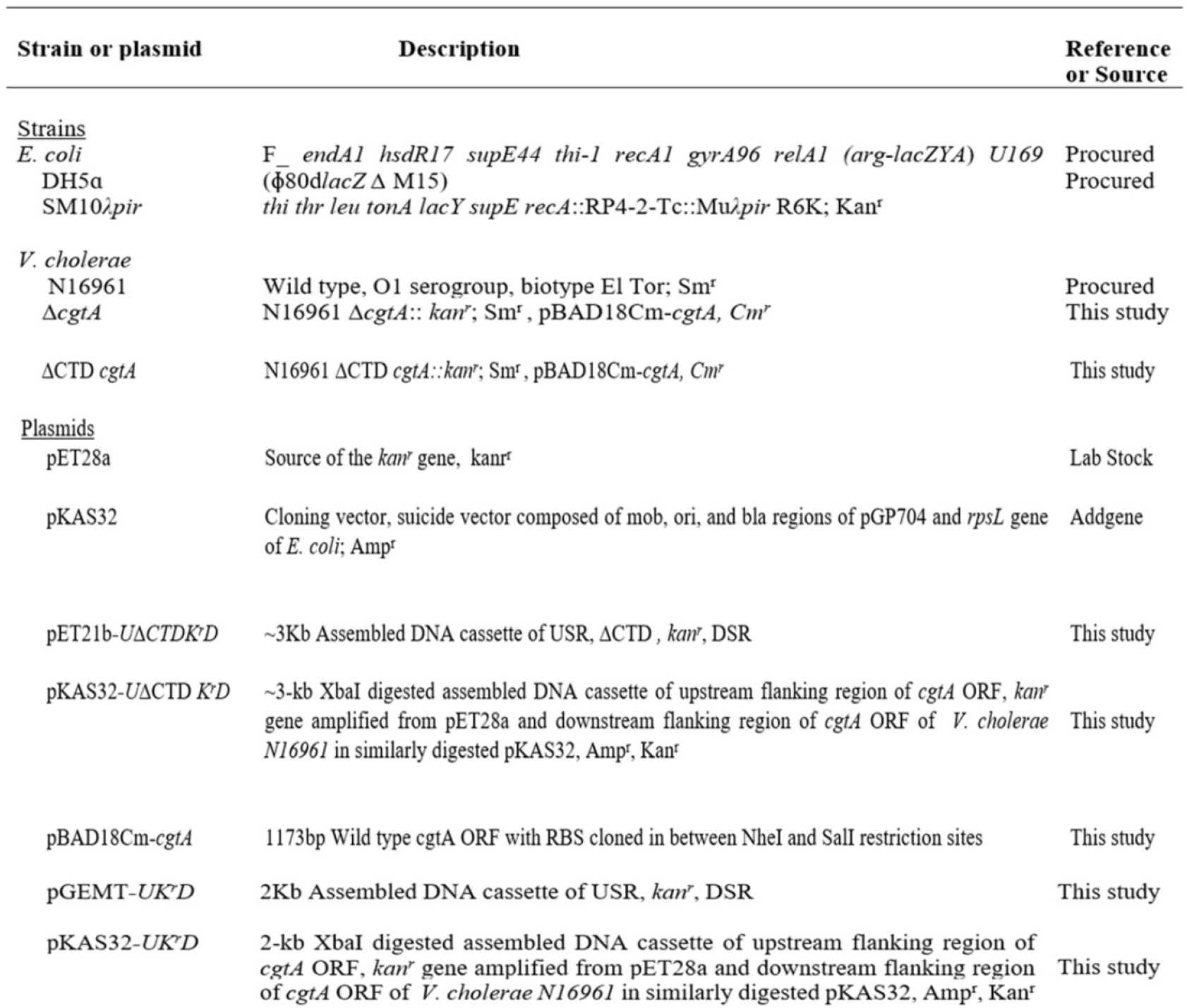
List of strains and plasmids used in this study

### 3.4 Complementation of the V. cholerae △cgtA::kan^r^ and △CTD cgtA::kan^r^ with wild-type CgtA expressed from pBADl8Cm vector

In order to delete *cgtA*, an essential gene, from the genome of *V. cholerae*, it has to be supplied *in trans* by an expression vector. In this case, CgtA_VC_ was expressed from arabinose inducible *araBAD* promoter from pBAD18-Cm expression vector. CgtA_VC_ open reading frame (ORF) was amplified (~1173bp) from the genomic DNA using primers 7 and 8. The primer 7 had NheI restriction site with a Ribosome Binding site (RBS) and the primer 8 had SalI restriction site. The amplified ORF was digested with NheI and SalI restriction enzymes, purified and ligated to a similarly digested pBAD18-Cm. The positive clones were screened by colony PCR, plasmid extraction, digestion, visualization of a 2Kb band after gel electrophoresis and DNA sequencing. The pBAD18Cm-cgtA was electroporated into *V. cholerae* cells and selected on LBA plate with chloramphenicol. Electrocompetent cells of *V. cholerae* were prepared by growing the cultures to mid-logarithmic phase, washing the cells with 2 mM Calcium chloride solution twice and resuspending the cells in 10% glycerol. Electroporation was performed in MicroPulser electroporation cuvette, 0.2 cm gap, according to the following conditions: C = 25 μF; PC = 200 ohm; V = 3.O kV

### 3.5. Verification of the expression of CgtA^VC^ from pBAD18-Cm expression vector

From an overnight culture of *V. cholerae*/pBAD18cm-*cgtA*, two subcultures were performed. One supplemented with 0.1% arabinose (test) and the other one without arabinose (control). After an induction of 5 h, the cells were harvested, lysed and checked on 12% SDS-PAGE stained with coomassie blue staining.

### 3.6. Growth studies of in vivo mutant strains of V. cholerae

For the study of the growth curves, each of the *V. cholerae* strains were sub-cultured in 100 ml of medium in 500 ml conical flasks from overnight growing pre-cultures after diluting to an optical density at 600 nm [OD_600_] of ~0.02 and analyzed in. During the incubation period, OD_600_ was taken at 30-min intervals using the EppendorfBioSpectrometer^®^ fluorescence instrument. The growth curves were performed in replicates. The data was plotted and analyzed statistically using GraphPad Prism 5.0software (GraphPad Software, San Diego, CA, USA).

### 3.7 Resuscitation of persister cells

The overnight primary cultures of all the *V. cholerae* strains in this study were grown without arabinose supplementation to stop the *in trans cgtA* expression (except for low level of leaky-expression as reported for pBAD18-Cm vector in some reports). The primary culture was washed twice in M9 minimal media and after 1:50 times dilution, the secondary culture in M9 minimal media was inoculated. The cultures were allowed to grow for 38 h (late stationary phase) at 37 °C on 200 rpm shaker. To check if any persister cells had formed under starvation conditions, resuscitation studies was carried out. Briefly, the persister cells were resuscitated by releasing the nutritional stress at 37°C. The plates were observed at various time-points of incubation. After overnight incubation, the cells were estimated by calculating the Log CFU/ml. Following this, the plates were further incubated till 24 h to check for the resuscitation of persister cell population.

### 3.8. Swarm Plate Assay

The soft agar plates were incubated at 37°C for 6 h followed by measuring the swarm diameter for each of the following *V. cholerae* N16961 strains: Wild-type, Δ*cgtA::kan^r^* and △*CTD cgtA::kan^r^*. LB soft-agar (1% Tryptone, 0.5% NaCl and 0.3% Agar) swarm plates were used to assess the chemotactic behavior. With the help of sterile toothpicks, soft-agar plates for swarming assay were inoculated by touching a bacterial colony on the LB agar plate and stabbing the toothpick to the soft-agar plate. The plates were incubated at 30°C for 6h under aerobic conditions. Plates were photographed with Syngene G: BOX imaging station. An average of three experiments was taken to calculate the swarming diameter followed by statistical analysis.

### 3.8. Scanning Electron Microscopy

All the strains of *V. cholerae* for this study were collected from exponential phase either in LB broth or M9 minimal media supplemented with casamino acids and glucose. The cells were harvested at 6000 RPM for 15 minutes. The cells were collected and washed with 1X PBS buffer twice. Then the cells were fixed in 2.5% glutaraldehyde prepared in 1X PBS for 2 hours. The sample was mixed gently by gentle pipetting in thirty minutes. This was followed by washing the cells twice with 1X PBS and serially dehydrating with various percentages of ethanol (from 30% −100%). The cells were then loaded onto broken glass coverslip and dried in vacuum. This was followed by coating with platinum. Samples were then observed under field emission scanning electron microscopy (FESEM) (SUPRA 55 VP-4132 CARL ZEISS).

### 3.9 Genome wide stress response induced by CgtA depletion by transcriptome analysis using RNA-Seq of V. cholerae strains

1μg of total RNA was taken for rRNA depletion using QIAseq FastSelect –5S/16S/23S Kit (Catalog: 335925, Invitrogen) according to manufacturer’s protocol. This method offers selective depletion of bacterial 5S/16S/23S rRNA. In brief, the RNA sample was hybridized to the FastSelect 5S/16S/23S probe followed by the removal of this complex through QIAseq Beads included in the kit. Thus, depleted RNA was eluted with NFW. NEBNext Ultra II RNA Library Prep Kit (Catalog: Catalog: E7775S, New England Biolabs) for Illumina was used for the library preparation. The enriched transcriptome was chemically fragmented in a magnesium-based buffer at 94°C for 15 minutes. The fragmented samples were primed with random hexamers, and reverse transcribed to form cDNA and the first strand cDNA reactions were converted to dsDNA. The double stranded cDNA fragments obtained were cleaned up by using AMPure beads (Catalog: A63881, Beckman Coulter). The cDNA fragments undergo end repair where in the mix converts the overhangs resulting from fragmentation into blunt ends. The 3’ to 5’ exonuclease activity of end repair mix removes the 3’ overhangs and polymerase activity fills in the 5’ overhangs. To the blunt ended fragments, adenylation was performed by adding single ‘A’ nucleotide to the 3’ ends. To the adenylated fragments, loop adapters (Platform Specific) are ligated and cleaved with uracil-specific excision reagent (USER) enzyme. The samples are further purified using AMPure beads (Catalog: A63881, Beckman Coulter). Furthermore, the DNA was amplified by 12 cycles of PCR with the addition of NEBNext Ultra II Q5 master mix, and “NEBNext^®^ Multiplex Oligos for Illumina” to facilitate multiplexing while sequencing. The amplified products were then purified using 0.9X AMPure XP beads (Catalog: A63881, Beckman Coulter) and the final DNA library was eluted in 15μl of 0.1X TE buffer. The library concentration was determined in a Qubit.3 Fluorometer (Catalog: Q33216, Life technologies using the Qubit dsDNA HS (High Sensitivity) Assay Kit (Catalog: Q32854, ThermoFisher Scientific). The Qubit^™^1X dsDNA HS Assay Kit. DNA HS Assay Kit contains High Sensitivity DNA reagent as well as buffers and two DNA standards. DNA HS assay reagent is one of the most sensitive detection dyes for the accurate quantitation of DNA/library in solution, with linear fluorescence detection in the range of 10pg/μl to 100ng/μl of DNA. Dye and the buffers were diluted at 1:200 ratio and 1μl of the library was mixed with the dye mix and incubated at RT for 2min and the readings were taken in the Qubit.3 Fluorometer (Catalog: Q33216, Life technologies). Prior to the sample’s measurement, the instrument was calibrated using the two standards provided in the kit. The library quality assessment was done using Agilent D1000 ScreenTape System (Catalog: 5067-5582, Agilent) in a 4150 TapeStation System (Catalog: G2992AA, Agilent) which is designed for analysing DNA molecules from 100 to 1000bp.1μl of the purified library was mixed with 3μl of D1000 sample buffer (Catalog: 5067-5583) and vortexed using IKA vortexer at 2000rpm for 1min and spun down to collect the sample to the bottom of the strip. The strip was then loaded on the Agilent 4150 TapeStation system. The sequence data was generated using Illumina HiSeq. Data quality was checked using FastQC and MultiQC software. The data was checked for base call quality distribution, % bases above Q20, Q30, %GC, and sequencing adapter contamination. All the samples have passed the QC threshold (Q20>95%). Raw sequence reads were processed to remove adapter sequences and low-quality bases using fastp. The QC passed reads were mapped onto indexed *Vibrio cholerae* reference genome (O1 biovar El Tor str. N16961) using Bowtie2 aligner. On average 98.55% of the reads aligned onto the reference genome. Gene level expression values were obtained as read counts using feature Counts software. Differential expression analysis was carried out using the DESeq2 package. Thread counts were normalized (variance stabilized normalized counts) using DESeq2and differential enrichment analysis was performed. Test sample was compared to the control sample. Genes with absolute log2 fold change ≥ 1 and p-value ≤ 0.05 were considered significant. The UpSetR R package was used to generate plots showing overlapping significant genes between conditions. The expression profile of differentially expressed genes across the samples is presented in volcano plots and heat map. The genes that showed significant differential expression were used for Gene Ontology (GO) and pathway enrichment analysis.

## 4. Discussion

The *V. cholerae cgtA* is essential for cell growth and the intracellular depletion of CgtA is deleterious for the viability of the cell. Attempts for in-frame deletion of *cgtA* from the *V. cholerae* chromosome were successful only in the presence of *in trans* supply of *cgtA* gene product. It was previously reported that the orthologues of *cgtA* is essential in *Escherichia coli* and *Neisseria gonorrhea. Raskin et al* and *Bhadra et al*., had also reported that *cgtA* is an essential gene in *V. cholerae*. But the domain specific essentiality of CgtA GTPase is reported in our work. Comparative morphological and physiological studies alongside whole genome transcriptome profiling between ΔC-terminal domain CgtA strain and CgtA depletion strain in our studies have provided valuable insights to understand the domain-wise essentiality as well as the function of CgtA GTPase in *V. cholerae*. We found that the C-terminal domain does not confer essentiality to the CgtA_VC_ GTPase. The viability of the ΔCTD CgtA strain is not affected unlike the CgtA depleted strain of *V. cholerae*. Thus, the essentiality of the CgtA GTPase lies in the NTD, GTPase domain or both. Since the NTD of CgtA takes part in the association with the 50S subunit of ribosome and thereby playing an essential role in ribosome biogenesis, the NTD is undeniably essential. Moreover, the signals for the instructions are conveyed to the conserved GTPase domain by the NTD via the flexible linker connecting the two domains. As a result of which the conformational changes are induced in the GTPase domain which either expedites GTP hydrolysis or the exchange of GDP for GTP. Hence, the GTPase domain is also an essential domain. The previous *in vitro* biochemical studies had shown that the CTD might have a regulatory role in the function of CgtA_VC_. To ascertain this, *in vivo* studies are necessary. The doubling time of ΔCTD CgtA strain is higher than the wild type strain. But, the doubling time of CgtA depleted strain was even many folds higher than the both the wild type and the ΔCTD CgtA strains of *V. cholerae*. Upon deleting the entire CTD of CgtA, pinhole sized colonies are observed. Hence, the Δ*CTD cgtA::kan^r^* is a small colony variant of the *V. cholerae* N16961 strain. When, complemented with wild type CgtA, the colony defect is not surpassed. Thus, the constitutive expression of the CTD truncated CgtA_VC_ is undesirable for the rescue of the growth defect by *in trans* expression of wild-type CgtA^VC^. The growth defect might be due to the constant bombardment of certain gene regulatory signals by the NTD and the GTPase domain leading to enhanced downstream effector functions which is impermissible for normal growth phenotype of the *V. cholerae* cells. In this case, the role of CTD becomes evident in regulating the function of first two domains of CgtA_VC_. Out of the three domains, only the sequence of the CTD is not conserved. Though, there are similarities in the conservation profile of amino acids in the gram-negative pathogens (enteric pathogens were taken into consideration in this work) and gram-positive pathogens separately. **(FIG. 12)** This might be a strong clue for the development of antibiotics against the CTD of CgtA_VC_ against various groups of phylogenetically closely related and distant human pathogens. Also, the CTD of CgtA is predicted to have an intrinsically disordered region (IDR), conferring flexibility to the protein which can further enhance its functionality (Anurag and Dash, 2009). Therefore, the outcome of this comparative study could be essential in curbing the problem of antibiotic resistance in addition to persistence in the pathogenic strains of *V. cholerae* considering CgtA to be an ideal drug target.

**FIG.12.**
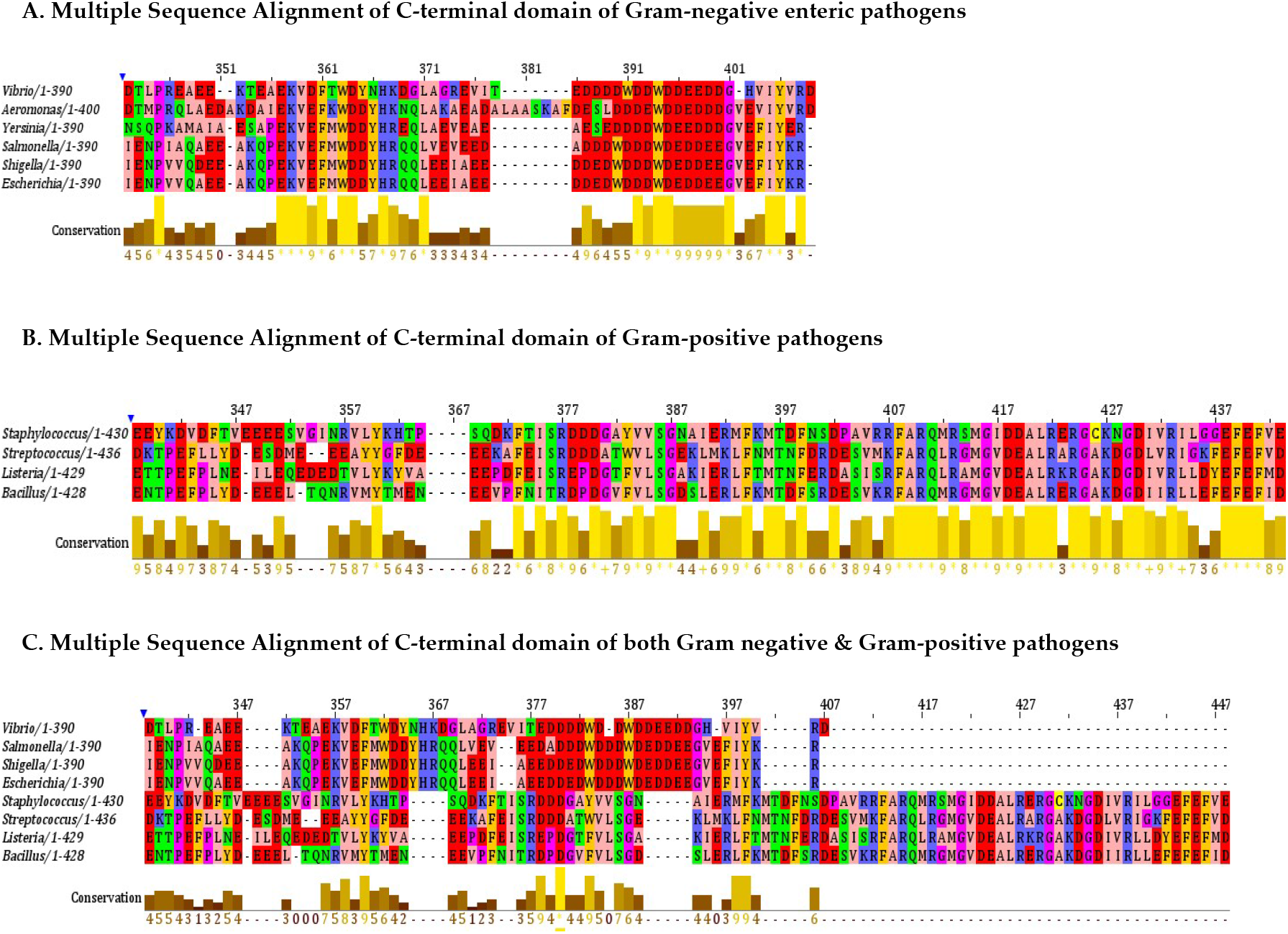
Multiple sequence Alignment (MSA) of the C-terminal domain of CgtA **(A.)** Gram-negative enteric pathogens, **(B.)** Gram-positive pathogens **(C.)** comparison between Gram negative and Gram-positive pathogens

Raskin *et al*. conducted the transcriptional profiling studies of CgtA depleted strain of *V. cholerae* in LB media and showed that CgtA is a regulator of the stringent response by comparing the *cgtA* depletion strain with the artificially induced stringent response in wild-type *V. cholerae*. We were interested to do a comparative transcriptome profiling between the Δ*CTD cgtA::kan^r^* and *cgtA* depletion strains to get a clear understanding of the global transcriptome landscape in the mutant strains as well as the wild-type strain. The RNA seq studies helped to define the domain specific regulatory activities. We grew all the strains in M9 minimal glucose media (casamino acids supplementation) to trigger the nutritional stress response in all the three strains. Apart from the genes co-regulated by the CgtA during the logarithmic phase of growth, the various other general stress response genes regulated by CgtA were also captured when grown in M9 minimal media, which would otherwise remain elusive when grown LB media. The regulatory activities of CgtA were predominantly carried out by the first two domains than the ΔCTD CgtA strain. While comparing the DEGs (<u>D</u>ifferentially <u>E</u>xpressed <u>G</u>enes) of the Δ*CTD cgtA::kan^r^* versus wild-type strain and Δ*cgtA::kan^r^* versus wild-type, the results from each of the categories clearly indicated that CTD has no role in transcriptional regulatory functions. Rather, the principal role of CTD is to coordinate the transcriptional regulatory activities of the first two domains of CgtA^VC^.

Nutritional stress in *Vibrio cholerae* is linked with persister cell phenotype. It is a mode of survival for *V. cholerae* under extreme environmental conditions. Serendipitously, we observed the microcolonies of CgtA depletion strain growing on nutrient rich LB media after plating the 38h old late stationary-phase culture and incubating for 24 hours. This clearly indicated the resuscitation of the persister cells that had formed due to the intracellular depletion of CgtA. While the wild-type and the ΔCTD CgtA strain had not shown any signs of persister cell resuscitation. Thus, the switch to form the persister cells is regulated by either of the first two domains of CgtA or both. Hence, it can be concluded that the CTD has no role in persister cell formation. We also observed that CgtA_VC_ controls the motility function of *V. cholerae* as well. The soft-agar motility assay. Apparently, the motility function is under the control of all the three domains of CgtA as the motility observed was highest for wild-type followed by gradual decrease in ΔCTD CgtA strain and lastly extremely reduced motility in case of CgtA depleted strain. It might be due to the fact that, the first two domains of CgtA preliminarily regulate the chemotactic genes, while the CTD aids in their regulatory functions. Deleting the CTD also reduces the motility but to a less extent than complete CgtA depletion from the cell.

Also, the CTD might act as a stress response sensor domain which senses and relays the nutritional stress signal to the N-terminal Obg and GTPase domain. Thus, the NTD and GTPase domain act as responders of nutritional stress. The manifestation of nutritional stress happens when CgtA is entirely depleted from the cell as evident from elongated *V. cholerae* cells observed under FESEM.

Thus, all of the findings indicate that the CTD serves as a nutritional stress response sensor which relays the signal to the other two domains. The CTD is not crucial for supporting cell’s viability but upon its deletion the growth of the *V. cholerae* is affected. Also, deletion of the CTD did not affect nutritional deprivation mediated persister cell formation, indicating that the CTD has no role in persistence. Further detail *in vivo* characterization of the three loops of the NTD will provide an insight of ribosome maturation by CgtA^VC^.

## Author Contributions

Experiments were designed by PPD and SD. Experiments were performed by SD. Experimental data were analyzed by PPD and SD. Manuscript was written by PPD and SD.

## Acknowledgement

Authors acknowledge Dr. Rupak Bhadra from the Indian Institute of Chemical Biology, Kolkata for providing the *Vibrio cholerae* N169661 strain and IISER Kolkata for proving the instrumentation facility. SD is supported by a PhD fellowship from IISER-Kolkata.

